# Label-Free Hemodynamic Mapping via Ultrafast Power Doppler-Enhanced Photoacoustic Imaging reveals early therapy-induced functional changes in the tumor

**DOI:** 10.1101/2025.11.19.688289

**Authors:** Deeksha M Sankepalle, Lucy Wei, Ronak Shethia, Pieter Kruzinga, Srivalleesha Mallidi

## Abstract

Real-time, non-invasive imaging of tumor vascular function is critical for assessing treatment response and informing therapeutic decisions. In this study, we present a dual-modality imaging approach that integrates Ultrafast power Doppler ultrasound (UPD) with multispectral photoacoustic (PA) imaging to quantify spatiotemporal changes in tumor oxygenation and perfusion following photodynamic therapy (PDT). Our results demonstrate that PDT induces dose- and time-dependent spatial heterogenic alterations in both oxygenation and perfusion. To better interpret the spatial interplay between tumor oxygenation and vascular perfusion, we developed a Vascular Function Index (VFI), which classifies tumor subregions into four functional quadrants based on perfusion and oxygenation status at every voxel within the 3D tumor region. This framework revealed dynamic, time-dependent changes in tumor hypoxia and perfusion, with high-dose treatment groups showing a progressive increase in hypoxic and non-perfused regions, underscoring the spatiotemporal nature of PDT-induced vascular response.

**Teaser:** Tumoral Blood flow and oxygenation interplay, assessed by label-free imaging, unravels heterogeneous therapy response.

## Introduction

Monitoring the early response of tumors to therapy is vital for evaluating treatment efficacy, optimizing therapeutic regimens, and preventing unnecessary interventions[1]. This is especially important in the case of aggressive solid tumors, which often possess abnormal and dysfunctional vasculature. Such disorganized vascular networks promote rapid tumor growth, facilitate metastasis, and significantly hinder the effective delivery of therapeutic agents and oxygen [2]. These vascular irregularities also contribute to the formation of hypoxic regions within the tumor microenvironment, making the tumors more resistant to many therapies, including chemotherapy and radiation [3]. As a result, surrogate functional markers of vascular function, such as microvascular density, perfusion, and oxygen saturation, are recognized as important early indicators of treatment response, particularly in therapies that target or affect the tumor vasculature [4].

There are several clinical vascular imaging techniques, such as Doppler ultrasound, Contrast-enhanced ultrasound, Computed Tomography (CT) Angiography, Magnetic Resonance (MRI) based techniques, and Optical techniques such as Optical Coherence Tomography angiography, etc. Ultrasound-based techniques such as Power Doppler have gained traction due to their non-ionizing nature and real-time imaging capabilities; however, their sensitivity is limited to vessels with relatively high flow rates [1, 5, 6]. A key limitation across these modalities is their reduced sensitivity to slow or intermittent flow within small-caliber vessels, which are often the most affected following vascular-targeted therapies. Signals from slow-moving erythrocytes in capillaries often fall below the noise threshold due to tissue motion [7]. As a result, these imaging platforms may fail to capture the early and most clinically relevant microvascular changes during treatment [8, 9]. Recent advances in ultrasound imaging, particularly in super-resolution imaging such as Ultrasound Localization Microscopy (ULM), have significantly improved the ability to visualize microvascular structures and detect slow-flowing blood within the tumor microenvironment [10]. These techniques offer spatial resolutions well beyond conventional ultrasound limits by tracking the movement of contrast microbubbles over time [11-15]. Despite the remarkable resolution, the dependency on contrast agents and lengthy acquisition times make these super-resolution techniques less practical for routine or longitudinal imaging, thereby driving interest in alternative label-free methods such as Ultrafast power Doppler ultrasound (UPD) [16, 17]. UPD ultrasound leverages ultrafast plane-wave imaging and advanced clutter filtering algorithms and significantly enhances sensitivity to imaging smaller blood vessels compared to conventional power Doppler methods [18-20].

Another aspect of the ultrasound-based imaging modalities, such as UPD, is the lack of ability to provide functional information of the vasculature, i.e., the blood oxygen saturation (StO_2_) levels. An imaging modality that gained immense popularity for assessing vascular function and StO_2_ is photoacoustic imaging (PAI). PAI is a hybrid technique that combines the high spatial resolution and deep penetration of ultrasound with the rich optical contrast of light absorption. By illuminating tissue with a nanosecond pulsed laser light and detecting the resulting ultrasound waves generated by thermoelastic expansion, PAI enables label-free visualization of vascular architecture and hemodynamics with high sensitivity. Importantly, the absorption spectra of hemoglobin and oxyhemoglobin are distinct, allowing PAI to directly quantify blood oxygen saturation and map hypoxic regions within tumors without requiring exogenous agents [21-25]. This functional dimension distinguishes PAI from most conventional vascular imaging modalities, making it particularly valuable in monitoring responses to anti-angiogenic or oxygen-dependent therapies[21, 23, 26].

PAI is inherently non-ionizing, cost-effective compared to CT or MRI, offers deeper penetration than other optical angiography techniques, and can be seamlessly integrated with existing ultrasound platforms, which accelerates its clinical adoption [21, 23, 27-29]. For example, Irisawa et al. demonstrated the superior sensitivity of PAI over Doppler ultrasound in detecting superficial blood vessels in humans [30]. Building on this, Zhao et al. reviewed conventional scoring techniques for evaluating rheumatoid arthritis and reported that PAI exhibited superior sensitivity in identifying vasculature compared to Doppler ultrasound [31]. Similarly, Peng et al. corroborated these findings, showing that PAI can detect inflammation and disease progression at earlier stages than conventional Doppler-based methods [32]. While PAI excels in functional imaging, its sensitivity to very slow flow dynamics and small-caliber vessels can still be challenged by background noise and limited temporal sampling. This is where synergistic approaches that combine modalities hold significant promise. Combining UPD with PAI enables a comprehensive, multi-parametric assessment of tumor vasculature by integrating structural, flow, and functional information. While UPD is highly sensitive to slow and microvascular blood flow [33], PAI complements it by providing hemoglobin and oxygenation measurements, together enabling early detection of therapy-induced vascular changes such as pruning, normalization, or hypoxia onset.

Recent advances in ultrasound imaging consoles have led to the acquisition of photoacoustic (PA) and UPD images simultaneously [34] in healthy organ systems such as the brain or kidney. Leveraging these advances, we optimized the acquisition parameters to monitor the tumor vasculature changes occurring (flow and function) due to a light-activatable therapy called photodynamic therapy (PDT), which offers spatial precision in inducing cytotoxicity while minimizing collateral damage to surrounding healthy tissues. In PDT, the therapeutic process begins with the administration of a photosensitizer (PS), which selectively accumulates in tumor tissue. Upon irradiation with light of a specific wavelength, the photosensitizer becomes excited and interacts with molecular oxygen present in the tissue, generating reactive oxygen species (ROS) [35]. These ROS mediate localized oxidative damage, leading to cell death and tissue destruction in the illuminated region [36, 37].

Importantly, PDT exerts its effects through a dual mechanism: direct cytotoxicity to tumor cells and disruption of the tumor vasculature [38]. Both pathways contribute to overall therapeutic efficacy, but vascular damage is particularly significant in the context of solid tumors. Several studies have leveraged PAI to non-invasively assess tumor oxygen saturation following PDT and other vascular-targeted therapies [22, 26, 39-41]. For instance, prior work by Mallidi et al. demonstrated that PAI could detect significant reductions in tumor oxygenation as early as 6 hours post-PDT [22]. However, StO_2_ alone may not fully capture the extent of vascular disruption caused by PDT. Given that ROS generated by the PS during light activation consumes local oxygen, decreases in oxygenation may reflect both therapeutic effect and photosensitizer activity rather along with structural vascular damage [42]. This limitation becomes more pronounced when assessing changes in the tumor microvasculature, where subtle alterations in flow and vessel integrity are more difficult to detect with oxygenation metrics alone. UPD offers a compelling, non-invasive solution to monitor these vascular changes in real time, enabling deeper insight into treatment efficacy [43, 44].

More importantly, to better gauge the spatial interplay between tumor oxygenation and vascular perfusion following PDT, we introduce a composite imaging metric named the Vascular Functional Index (VFI). This is possible due to the acquisition of the flow and StO_2_ information within the same region of interest in the tumor simultaneously at multiple time points post-treatment. In this study, we demonstrate that the VFI maps showcase the heterogeneity in the temporal dynamics of vascular disruption and therefore can better identify early markers of sub-optimal treatment efficacy.

## Results

### Validation of dual modality images with histology

To validate the spatial and functional accuracy of our dual-modality imaging approach, Figure 1 presents a direct comparison between *in vivo* HbT and StO_2_ maps and *ex vivo* histological analyses, Hematoxylin and Eosin (H&E) staining, and immunofluorescence (IF) labeling for vascular (CD31) and hypoxia (pimonidazole) markers. Three representative mice are shown to illustrate the tumor vascularization and oxygenation profiles captured by the imaging system. In Mouse 1 (Fig. 1, top row), the HbT and StO_2_ maps reveal a well-vascularized and oxygenated tumor, evidenced by high HbT values and red regions in the StO_2_ map. Corresponding IF images show widespread CD31 expression, with pimonidazole staining largely restricted to the tumor periphery or surrounding connective tissue. These peripheral regions, highlighted by yellow boxes, show low StO_2_ and positive hypoxia staining, indicating insufficient oxygen delivery despite vascular presence. In Mice 2 and 3, more heterogeneous vascular and oxygenation patterns are observed. Yellow ROIs in both animals demonstrate regions of low StO_2_ that spatially co-localize with pimonidazole staining, consistent with functionally hypoxic vasculature. These areas retain elevated HbT values, suggesting the presence of blood volume without effective oxygen exchange evidently seen in the CD31 stain as well. In contrast, white ROIs across all samples highlight regions with high StO_2_ and absent pimonidazole staining, consistent with well-perfused, normoxic tissue. Collectively, these findings validate the ability of PAI to capture spatially resolved functional oxygenation and to distinguish hypoxic from normoxic tumor regions.

**Fig. 1:**
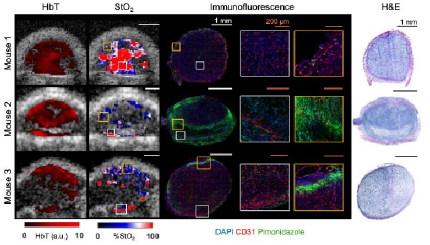
Validation of photoacoustic images for assessment of tumor vascularization and hypoxia. *Left panels*: Total hemoglobin (HbT) and oxygen saturation (StO_2_) maps in tumor cross-sections from three different samples (rows). *Middle panels*: Corresponding immunofluorescence images highlighting vascular and hypoxia markers. DAPI (nuclei, blue), CD31 (endothelial cells, red), and pimonidazole (hypoxia, green). Yellow (regions with hypoxia) and white boxes indicate regions of interest shown at higher magnification (right of each sample). *Right panels*: H&E stained sections. Scale bars: 1 mm (white, whole tissue), 200 µm (orange, zoomed-in regions).

Similarly, Figure 2 demonstrates the functional accuracy of UPD imaging by comparing it with IF images stained for CD31 (red, vascular structure) and tomato lectin (green, perfusion marker). Figures 2A and 2B show the PDI and HbT maps for a representative tumor. Figure 2C presents a top-view photograph of the tumor. The dashed line indicates the imaging plane, and the blue arrow points to superficial skin vessels that correspond to prominent Doppler signals in Fig. 2A.

**Fig. 2:**
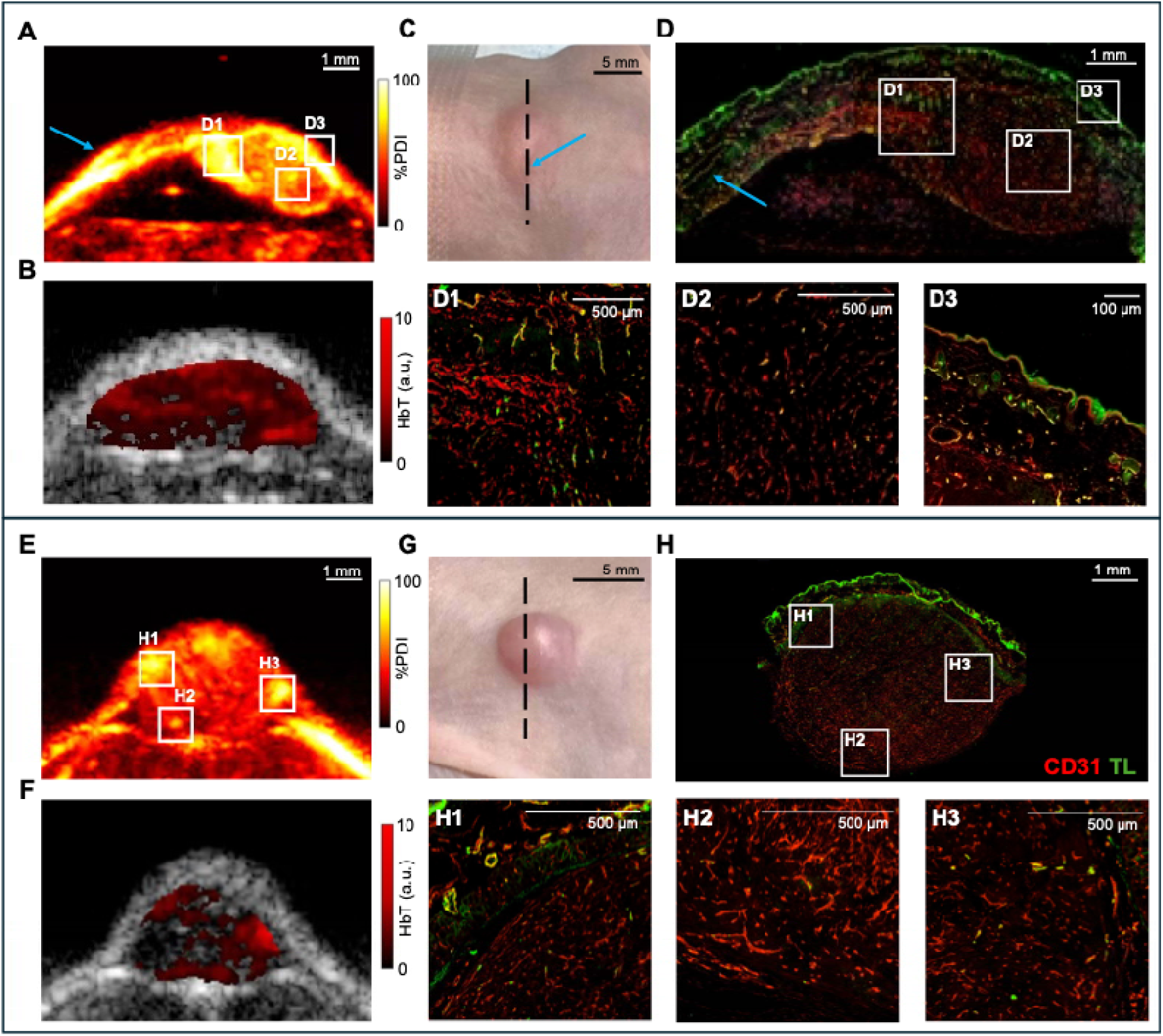
Strong correlation between *in vivo* UPD images and *ex vivo* histological assessment of tumor vascularization and permeability. **A, E)** Power Doppler intensity (%PDI) maps highlighting areas of high vascular permeability in two representative tumors (white boxes indicate regions of interest; dashed lines show the imaging plane). **B, F)** Corresponding hemoglobin concentration (HbT) maps overlaid on grayscale US images, showing elevated vascular content in the same regions. ***C, G)*** Photographs of the tumors with dashed lines indicating imaging planes. ***D, H)*** Immunofluorescence sections showing vasculature (CD31, red) and perfusion marker tomato lectin (TL, green). The boxed regions (D1–D3, H1–H3) correspond to high-resolution insets that reveal detailed vascular structure and tracer localization. High concordance is observed between elevated %PDI in imaging and areas of strong tracer signal in histology, validating the ability of photoacoustic imaging to non-invasively assess tumor vascular function and permeability. Scale bars: 1 mm (whole tissue), 500 µm or 100 µm (insets).

Figure 2D shows the corresponding IF image of the imaged tumor cross-section. Specifically, inset D1 corresponds to areas with a strong Doppler signal, and the IF images show dense lectin-positive staining, indicating well-perfused blood vessels. In contrast, region D2 demonstrates low Doppler signal intensity and corresponds to sparse green staining on the IF image, suggesting limited or absent perfusion despite the presence of vasculature. Region D3, located in the surrounding skin, serves as a control area and confirms consistent perfusion, as expected in non-tumor tissue.

Similarly, Figures 2E–H present a second tumor to illustrate vascular heterogeneity. Inset H1 highlights a peripheral region of the tumor with a high Doppler signal, validated by the presence of both a large vessel just outside the tumor boundary and several smaller perfused vessels beneath it, as shown by green lectin staining. Inset H2 corresponds to a region with lower Doppler intensity, which is validated by a relative paucity of perfused vessels on the IF image. Inset H3, located within the tumor core, again shows a strong Doppler signal and a dense pattern of lectin-positive vessels.

The HbT maps exhibit strong spatial concordance with CD31 in both tumors, confirming the structural vascular information captured by PAI. However, not all CD31-positive vessels are perfusive, as highlighted by discrepancies between structural and Doppler images. Perfused vessels (identified by the lectin stain in green) align well with areas of high Doppler intensity in PDI maps. Together, these data validate the ability of UPD to noninvasively assess spatial heterogeneity in tumor perfusion with high specificity to functional vasculature.

### Dose-dependent changes in StO_2_ and PDI

The integration of PA imaging with UPD ultrasound enabled volumetric imaging of tumor oxygenation and vascular dynamics in subcutaneous glioma xenografts. This combined imaging approach provided complementary insights into the functional and structural changes in tumor vasculature following PDT. As observed in our previous study [22], PDT induced a marked reduction in tumor oxygen saturation, particularly evident at 6 hours post-treatment in the 100 J/cm^2^ group.

Pre-treatment imaging revealed well-oxygenated and densely vascularized tumors, as seen in both the PA and Doppler images (Fig. 3A). Following 100 J/cm^2^ irradiation, a substantial decrease in StO_2_ was detected at 6 hours, reflecting acute hypoxia likely due to the consumption of oxygen during ROS generation and vascular shutdown [45]. By 24 hours post-PDT, partial reoxygenation was observed in a few areas in PA images, consistent with prior findings [22, 46]. UPD images showed a continued absence of detectable microvascular flow, indicating poor perfusion despite apparent increases in oxygen levels.

**Fig. 3:**
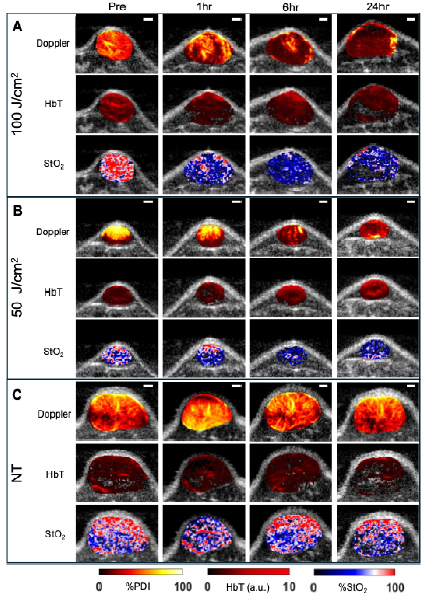
Dual-modality imaging reveals dynamic vascular and oxygenation changes following PDT in various groups. **A**)100 J/cm^2^ - Pre-treatment HbT, StO_2_ and Doppler images show well-oxygenated and vascularized tumors. At 6 hours post-PDT a reduction in StO_2_ indicates acute hypoxia, accompanied by loss of Doppler-detectable flow. Partial reoxygenation is observed at 24 hours in PA maps, though microvascular flow remains absent, suggesting limited vascular recovery. **B)** A low light dose group (50 J/cm^2^) shows similar temporal trends of transient oxygen depletion along with reduction in Doppler signal from vascular damage. **C)** No-treatment (NT) controls (PBS + 100 J/cm^2^ light) maintain stable StO_2_ and vascular profiles throughout, confirming PDT-specific effects.

To further evaluate the sensitivity of the dual-modality imaging strategy in detecting suboptimal PDT effects, an additional treatment group was included that received a lower light dose at 50 J/cm^2^, while maintaining the same liposomal BPD concentration. This group exhibited similar temporal trends (Fig. 3B), an initial drop in StO_2_ and Doppler signal followed by partial oxygenation recovery in a few tumor regions without corresponding microvascular reappearance. In contrast, the no-treatment (NT) control group, which received a sham PBS injection and 100 J/cm^2^ light exposure, exhibited stable oxygenation and vascular profiles over the 24-hour period (Fig. 3C), confirming the specificity of the observed changes to PDT-mediated effects.

To further evaluate the effects of PDT on tumor oxygenation and perfusion, group-wise quantitative analyses of three-dimensional average StO_2_ and PDI values were performed and are summarized in Figure 4. These average values were calculated by analyzing the central 80% of each tumor volume, intentionally excluding peripheral regions to account for potential inconsistencies in light distribution during PDT. To minimize inter-subject variability and highlight relative treatment-induced changes, all StO_2_ and PDI values were normalized to each animal’s pre-treatment baseline. Statistical analysis was conducted using two-way ANOVA with Tukey’s post hoc test. There were no significant differences in tumor volume among the groups at baseline, confirming effective randomization and consistent baseline tumor physiology.

**Fig. 4:**
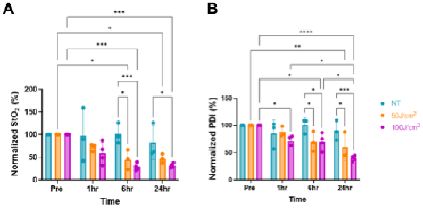
Quantitative assessment of dose dependent of changes in StO_2_ and PDI across treatment cohorts and time points. **A**) Baseline-normalized StO_2_ values across the NT (n=3), 50 J/cm^2^ (n=3), and 100 J/cm^2^ (n=4) PDT groups show no statistically significant differences at pre-treatment time points, indicating consistent physiological baselines. **B)** Power Doppler intensity (% PDI) revealed a sustained decrease beginning at 1 hour in the 100 J/cm^2^ group, while the 50 J/cm^2^ group exhibited a significant reduction only at 24 hours. Both StO_2_ and PDI responses exhibited fluence- and time-dependent trends. The 100 J/cm^2^ group showed a significant drop in StO_2_ as early as 6 hours, persisting to 24 hours, whereas the 50 J/cm^2^ group showed a delayed but significant reduction at 24 hours. The NT group showed no significant changes over time in either parameter, confirming treatment specificity. Data are presented as mean ± SEM. Statistical comparisons were performed using two-way ANOVA with Tukey’s post hoc test;

Fluence-dependent efficacy of PDT has been well documented in previous studies [22, 47-49], particularly in relation to the timing and magnitude of oxygenation and vascular changes. In line with this literature, our findings demonstrate that both StO_2_ and PDI responses are temporally and fluence dependent. In the 50 J/cm^2^ treatment group, oxygen saturation showed a statistically significant decline at both 6 and 24 hours post-treatment. The 100 J/cm^2^ group exhibited a similar pattern, with a more pronounced and earlier decline observed at 6 hours, persisting through 24 hours. Although localized areas of reoxygenation were observed in some tumors at 24 hours, the overall StO_2_ increase relative to 6 hours did not reach statistical significance.

Similarly, analysis of PDI revealed that the NT group maintained stable perfusion across all time points, consistent with the absence of therapeutic intervention. The 50 J/cm^2^ group showed no statistically significant changes at early time points (1 and 6 hours), though a significant decrease in PDI was evident by 24 hours, suggesting a delayed vascular response at this fluence. In the 100 J/cm^2^ group, PDI exhibited a significant reduction as early as 1 hour post-treatment, with a continued decline that persisted through 6 and 24 hours. This early decline in perfusion is clearly visualized in the PDI maps (Fig. 3A). These results suggest that higher PDT fluence induces more rapid and sustained vascular and oxygenation disruption, underscoring the importance of dose optimization for maximizing therapeutic efficacy.

### Combined VFI parameter reveals early treatment response

Building upon the earlier observation of significant early vascular changes in the 100 J/cm^2^ group, as captured by UPD imaging, we utilized the VFI maps to further explore the spatial and temporal correlation between tumor oxygenation and perfusion. VFI map provides a pixel-wise classification of tumor regions into four functional quadrants based on predefined thresholds for StO_2_ and PDI: Q1 (high StO_2_, high PDI – red), Q2 (high StO_2_, low PDI – green), Q3 (low StO_2_, high PDI – blue), and Q4 (low StO_2_, low PDI – white) as shown in Fig. 5A. These thresholds were empirically determined for this specific tumor model but may require adaptation for different tumor phenotypes or experimental conditions.

**Fig. 5:**
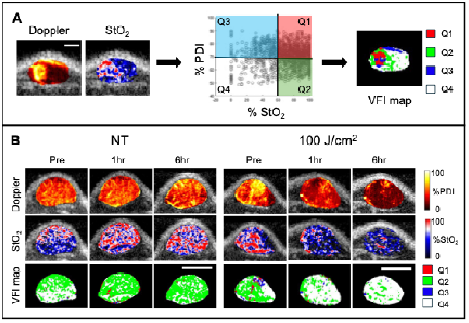
Vascular Function Index (VFI) mapping reveals spatiotemporal changes in tumor oxygenation and perfusion following PDT. **A)** Schematic of the VFI classification map based on pixel-wise thresholds for StO_2_ and PDI, dividing pixels into four functional quadrants: Q1 (high StO_2_, high PDI – red), Q2 (high StO_2_, low PDI – green), Q3 (low StO_2_, high PDI – blue), and Q4 (low StO_2_, low PDI – white). **B)** Representative VFI maps from the NT and 100 J/cm^2^ groups over time. In the treated group, a progressive decline in Q1 and Q2 regions was observed, consistent with PDT-induced impairment in vascular function and oxygen delivery. Q4 regions became increasingly dominant at 24 hours, indicating extensive vascular shutdown and hypoxia. In contrast, the NT group showed stable VFI profiles across time points, confirming vascular and oxygenation stability in the absence of treatment.

Representative VFI maps for the NT and 100 J/cm^2^ groups are shown in Fig. 5B. Consistent with biological expectations, Q3 regions (low oxygenation with high perfusion) were rare, likely due to the limited physiological plausibility of active blood flow with poorly oxygenated regions [42, 50]. A more prominent observation was the progressive decline in Q1 (well-perfused, oxygenated regions) and Q2 (well oxygenated but low perfused regions) over time in the 100 J/cm^2^ group, as shown in red and green in Fig. 5B. This suggests a gradual loss of vascular function and oxygen delivery post-PDT. Conversely, the NT group exhibited minimal changes across all VFI quadrants, reinforcing the stability of vascular function in the absence of treatment. Notably, Q4 (low oxygenation and low perfusion) regions became increasingly dominant in the 100 J/cm^2^ group at later time points, indicating the expansion of non-functional vasculature.

To quantitatively evaluate spatial changes in vascular function, the proportion of pixels corresponding to each VFI quadrant was calculated by determining the ratio of quadrant-specific pixels to the total number of tumor pixels in each imaging frame. This allowed for a pixel-wise assessment of regional vascular dynamics. Figure 6A displays donut plots representing the relative distribution of all four VFI parameters, VFI_Q1, VFI_Q2, VFI_Q3 and VFI_Q4 across treatment groups and time points. To better visualize treatment-induced changes, all values were normalized to their respective pre-treatment baselines. This approach enables a comparative, intuitive representation of how each functional quadrant evolves over time post-PDT. As anticipated, VFI_Q1 regions declined progressively in PDT-treated groups, reflecting vascular damage and reduced oxygen delivery. Given the consistently low fractions of VFI_Q1 and VFI_Q3 compared to VFI_Q2 and VFI_Q4, further analysis focuses primarily on VFI_Q2 and VFI_Q4, which capture the most prominent functional shifts post-treatment.

**Fig. 6:**
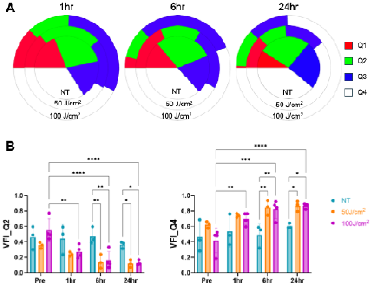
Vascular Function Index (VFI) reveals spatiotemporal disruption of tumor vasculature post-PDT. **A**) Donut plots illustrating the normalized fractional distribution of tumor regions into four quadrants, VFI_Q1 (high StO_2_, high PDI, red), VFI_Q2 (low StO_2_, high PDI, green), VFI_Q3 (high StO_2_, low PDI, blue), and VFI_Q4 (low StO_2_, low PDI, white) at 1 hour, 6 hours, and 24 hours post-treatment in NT, 50 J/cm^2^, and 100 J/cm^2^ groups. A progressive increase in Q4 regions and declines in Q1, Q2 and Q3 regions was observed in treated groups over time, especially at 100 J/cm^2^, indicating loss of vascular function and oxygen delivery. **B**) Quantitative analysis of VFI_Q2 and VFI_Q4 fractions over time. The 100 J/cm^2^ group demonstrated a more pronounced temporal trend, indicating fluence-dependent deterioration of low-perfused but normoxic vasculature. Data are represented as mean ± SEM; *p < 0.05, **p < 0.01, ***p < 0.001, ****p < 0.0001 (two-way ANOVA with Tukey’s post hoc test).

Statistical analysis of VFI fractions (Fig. 6B) demonstrated a significant increase in VFI_Q4 and a corresponding decrease in VFI_Q2 in both PDT-treated groups (50 J/cm^2^ and 100 J/cm^2^) compared to the NT group at 6 and 24 hours post-treatment (p < 0.05). These findings underscore the effectiveness of the VFI framework in capturing spatially resolved, localized deterioration of vascular function because of PDT. Notably, the 100 J/cm^2^ group exhibited a pronounced and statistically significant temporal trend, with progressive increases in VFI_Q4 and concurrent decreases in VFI_Q2 from pretreatment to 24 hours post-treatment. This suggests a dose-dependent acceleration in the collapse of functionally poorly perfused but well oxygenated regions, likely reflecting more extensive vascular shutdown and oxygen depletion at higher fluence levels.

The VFI maps provide valuable insight into how different tumor regions respond to treatment over time. Tumors are inherently heterogeneous in both structure and function, and this heterogeneity significantly influences therapeutic outcomes. VFI maps can identify regions that are well-perfused and oxygenated, which may remain viable longer post-treatment, as well as areas that become rapidly hypoxic and poorly perfused, indicating effective vascular shutdown. Importantly, regions exhibiting high oxygenation, but poor blood flow (Q2, green), may serve as early indicators of incomplete treatment response. Understanding this spatial variability allows for more precise monitoring and stratification of tumor response [51]. It also opens the door to not only PDT but other treatments and treatment personalization, allowing clinicians to potentially adjust therapeutic parameters such as light fluence, drug dosage, or timing of intervention based on the functional characteristics revealed by VFI.

## Discussion and conclusion

The combination of UPD and multispectral PAI provides a label-free, non-invasive means to longitudinally monitor tumor physiology *in vivo*. Consistent with previous reports [22, 52], our results demonstrate that PA-derived oxygen saturation changes are dose- and time-dependent and serve as a reliable biomarker for PDT response. Furthermore, the addition of UPD imaging reveals alterations in microvascular flow, which, when used alongside StO_2_ maps, elucidates the interplay between oxygen supply and blood flow post-treatment. The early decline in PDI, observed as soon as 1 hour post-treatment in the 100 J/cm^2^ group, and the corresponding expansion of Q4 (low oxygenation and low perfusion) regions in the VFI map, underscore the potential of this dual-modality system in detecting early treatment-induced vascular shutdown.

Despite these promising findings, certain hardware limitations remain. While broadband transducers effectively capture high-spatial-resolution PA images, Doppler imaging benefits from narrowband transducers optimized for detecting blood flow. Thus, the development of dual-range or hybrid transducer systems that combine both broadband and narrowband functionalities could be pivotal in fully realizing the potential of integrated PA and UPD imaging for functional tumor assessment. Furthermore, improving the sensitivity and functional accuracy of UPD imaging will rely heavily on advanced filtering and signal processing techniques, such as Acoustic Sub-Aperture Processing (ASAP)[53], Spatiotemporal Non-Local Means Filtering [54], and Spatial Angular Coherence Factor weighting [55]. Integration of these methods into the UPD pipeline can substantially improve the detection of slow and microvascular flow, particularly in deeper tumor regions where Doppler sensitivity is compromised due to attenuation from the use of high-frequency transducers. These enhancements are expected to improve vessel conspicuity without sacrificing frame rate, allowing more accurate delineation of perfused vasculature within heterogeneous tumor environments.

Looking forward, the inclusion of microbubble- or nanobubble-based contrast-enhanced ultrasound techniques could further enhance vascular imaging sensitivity, particularly in detecting subtle flow within the tumor. These imaging modalities, when used in conjunction with the existing PA and UPD framework, may provide an even more comprehensive understanding of vascular integrity, permeability, and response to therapy. In addition, the PA component, known to accurately quantify blood oxygenation, could be further leveraged to inform on the hypoxia-to-normoxia transitions that occur during tumor reoxygenation, a process with known implications for tumor regrowth and treatment resistance. Moreover, targeted nanodroplets [41, 56] could allow simultaneous assessment of molecular expression and physiological function. Combined with longitudinal imaging using VFI, this approach may help not only in predicting therapeutic efficacy but also in dynamically guiding secondary interventions before tumor relapse occurs. The VFI metric enhances our interpretive capabilities by capturing heterogeneity in tumor response. For instance, the distinction between Q2 (High oxygenation and low perfusion) and Q4 regions can inform on the presence of transient versus permanent vascular damage, which may have implications for therapy planning and re-treatment timing. Moreover, the VFI framework offers a platform-independent method to map and quantify functional tumor subregions, which can be extended to other therapeutic modalities beyond PDT, including radiation and anti-angiogenic therapies. Together, these technologies represent a significant step forward in functional tumor imaging, with potential applications in both preclinical research and, ultimately, clinical translation for personalized therapy guidance.

## Materials and Methods

### Animal preparation and imaging protocol

A six-week-old male Foxn1nu nude mouse (The Jackson Laboratory) was used to establish the tumor model. Glioblastoma U87 cells were maintained in suspension culture with media containing 10% fetal bovine serum and 1% penicillin-streptomycin (100 U/mL), incubated at 37°C with 5% CO□. These cells (3 × 10□) were then suspended in 100 μL of a 1:1 mixture of Matrigel® and phosphate-buffered saline and were injected subcutaneously using a 1-mL tuberculin syringe fitted with a 28-gauge needle. Once the tumor reached a volume of 60–120 mm^3^, the mouse was anesthetized with isoflurane gas (2–3% for induction, 1.5% for maintenance) and placed on a temperature-controlled heating pad to preserve core body temperature. Respiration was continuously monitored during imaging to ensure stable physiological conditions. Acoustic coupling was facilitated with a layer of bubble-free, clear ultrasound gel applied directly to the tumor surface. Following the final imaging session at the 24-hour time point, mice were administered pimonidazole (60 mg/kg; Hypoxyprobe, Inc.) via tail vein injection to assess tumor hypoxia. Five minutes before euthanasia, Tomato Lectin (100 µg; Vector Laboratories) was injected intravenously to label therapy-induced vascular perfusion changes. After euthanasia, the tumors were extracted for further histological analysis.

### PDT treatment

The liposomal formulation of benzoporphyrin derivative (BPD) was synthesized using a thin film hydration and extrusion method, as previously described in Mallidi et al and Langley et al. [22, 57]. Briefly, the liposomes were synthesized using a thin film hydration and extrusion method. The lipids cholesterol (ovine) (20 μmol), 1,2-dipalmitoyl-sn-glycero-3-phosphocholine (2.5 μmol), 1,2-dioleoyl-3-trimethylammonium-propane (chloride salt) (1 μmol), and 1,2-distearoyl-sn-glycero-3-phosphoethanolamine-N-{methoxy (polyethylene glycol)-2000} (ammonium salt) (10 μmol) (Avanti Polar Lipids) along with BPD (MedChem Express) (0.25 μmol) in chloroform (Thermo Fisher Scientific) were added to a 10 ml round bottom flask. The chloroform solution was evaporated under a gentle stream of nitrogen. Once the film was completely dry, it was hydrated with 1 mL of Dulbecco’s PBS (Thermo Fisher Scientific), vortexed for 5 seconds, and then left in the hot water bath at 45°C for 10 minutes. The liposomal BPD solution was then vortexed vigorously and put on ice for 10 minutes. The freeze-thaw cycles were repeated for a total of 5 cycles. After the last freeze-thaw cycle, the solution was extruded (6 cycles) at 50°C through a 0.1 mm polycarbonate membrane (Cytiva Life Sciences).

Mice bearing U87 glioma tumors were randomly assigned to three treatment groups: no treatment (NT), low-dose PDT (50 J/cm^2^), and high-dose PDT (100 J/cm^2^). The NT group received an intravenous injection of 100 μL phosphate-buffered saline (PBS) followed by laser irradiation at 100 J/cm^2^ to serve as a control for light-only exposure. The PDT-treated groups received liposomal BPD at a dose of 0.5 mg/kg BPD equivalent via tail vein injection. Tumors were irradiated using a 690 nm continuous-wave laser (Modulight Corporation), with the laser beam delivered through a fiber-coupled setup post-drug light interval of 90 minutes (Irradiance was 100 mW at the tumor surface). The 50 J/cm^2^ and 100 J/cm^2^ received 500 seconds and 1000 seconds of irradiation, respectively.

### UPD and PAI system

All imaging was performed using a custom integrated ultrasound and PAI system comprising a Vantage 256 research ultrasound platform (Verasonics, Redmond, WA, USA) and a tunable optical parametric oscillator (OPO) laser system (Phocus HE Mobile, OPOTEK, Carlsbad, CA, USA). The laser was configured to operate at a pulse repetition frequency (PRF) of 10 Hz, with a tunable wavelength range from 690 to 950 nm [58]. For optical excitation, a fiber bundle integrated with the linear ultrasound transducer was coupled directly to the laser’s output port, providing uniform and efficient light delivery to the tissue surface.

After the animals were anesthetized and physiological stability was confirmed, PA images were acquired at 750 nm and 850 nm, wavelengths that facilitate differentiating oxygenated and deoxygenated hemoglobin for StO_2_ mapping. Following PA, the UPD ultrasound was acquired. The probe transmitted plane waves using 3-cycle pulses at a center frequency of 15.6 MHz. Each acquisition sequence consisted of 16 plane wave transmissions at steering angles uniformly distributed between -9 degrees and +9 degrees. The PRF was set to 5000 Hz. In total, 3600 angle-compounded frames were acquired per recording. Data was recorded continuously for 5.76 seconds. For each imaging session, 600 repetitions of ensemble were collected per buffer frame, and six such buffer frames were acquired sequentially. To enhance the signal-to-noise ratio and improve sensitivity to low-velocity flow, the resulting data were averaged to compute a single Power Doppler Intensity (PDI) image. For volumetric imaging of the tumor, the transducer was mounted on a computer-controlled precision linear motorized stage (X-LSM, Zaber Technologies), which enabled stepwise scanning across the tumor in the elevational direction. This setup facilitated full-volume acquisition of both Doppler and PA datasets across the tumor region. All raw channel data of UPD and beamformed B-mode ultrasound data were saved for subsequent offline processing and quantitative analysis. After each imaging session, the mouse was returned to its cage and monitored for full recovery. The imaging protocol described above was consistently applied at multiple time points: pre-treatment, and at 1 hour, 6, and 24 hours post-treatment to evaluate short-term vascular and oxygenation responses as shown in Fig. 7.

**Fig. 7:**
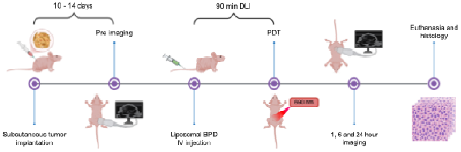
Experimental workflow for assessing photodynamic therapy (PDT) response using PA and Doppler imaging in a murine tumor model. Subcutaneous tumors were established and imaged pre-treatment to obtain baseline vascular and oxygenation maps. Mice received IV liposomal BPD, followed by a 90-minute drug light interval and 690 nm laser irradiation. Post-treatment imaging was performed to track PDT-induced changes. Tumors were then harvested for histological and immunofluorescence validation.

### Oxygen saturation and power Doppler image processing

All *in vivo* data processing was performed using MATLAB. As described in Fig. 8A, initially, raw PA images acquired at 750 nm and 850 nm were pre-processed by applying a threshold (calculated from an average of 3 mice at 750 and 850 nm wavelength illumination) to eliminate background noise. Following this, the images were spectrally unmixed to separate distinct chromophore contributions with our previously established unmixing algorithm [59]. Based on the ultrasound images, the tumor regions were identified, segmented, and subsequently overlaid on the unmixed images to visualize localized chromophore distributions. A 5 × 5 median filter (medfilt2) was then applied to the data, ensuring noise reduction and spatial smoothing of the StO_2_ map. Finally, a total hemoglobin (HbT) threshold was used to further suppress signals from areas with no hemoglobin presence (e.g., outside of the tumor region, such as from ultrasound gel), ensuring that the final StO_2_ measurements were confined to regions with meaningful vascular content.

**Fig. 8:**
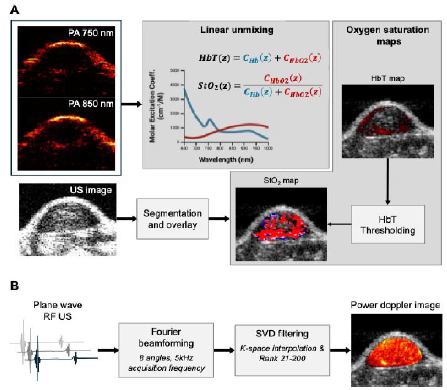
Processing pipeline for oxygen saturation and vascular function mapping using photoacoustic (PA) and ultrasound (US) imaging. **A**) Multi-wavelength PA images (750 and 850 nm) were acquired along with US. Linear spectral unmixing was applied to PA images to derive total hemoglobin (HbT) and oxygen saturation (StO_2_) maps. Co-registered US images enabled tumor segmentation and spatial overlay. Regions with physiologically relevant HbT levels were retained using thresholding to generate accurate oxygenation maps. **B**) Plane wave RF US signals were reconstructed using Fourier beamforming (8 angles, 5 kHz acquisition rate), followed by SVD-based clutter filtering (rank 21–200, K-space interpolation) to generate the power Doppler image.

Following the acquisition, the raw radiofrequency (RF) data were exported for offline processing. Beamforming was performed using ‘echoframes’, an open-source, real-time beamforming and power Doppler imaging library specifically developed for plane-wave ultrasound imaging. All acquisition parameters were input into the application to accurately reconstruct the beamformed data using a Fourier-domain beamforming algorithm optimized for UPD imaging [60]. The resulting beamformed data were subsequently interpolated by a factor of 200 in k-space. To isolate blood flow signals and suppress tissue clutter, a singular value decomposition (SVD) filtering approach was employed as described in Demene et al. [61]. Based on prior calibration studies, the threshold ranks for the SVD filter were empirically determined and fixed between 21 and 200. These values were found to provide optimal separation of slow-moving blood flow from stationary or slowly varying tissue components, resulting in high-quality PDI maps that accurately reflect tumor microvascular perfusion, as shown in Fig. 8B.

The VFI map, a pixel-wise, spatially resolved parameter, integrates StO_2_ maps with PDI data to classify each voxel within the tumor into one of four distinct functional quadrants:

- Q1 (Red) – High oxygenation and high perfusion
- Q2 (Green) – High oxygenation and low perfusion
- Q3 (Blue) – Low oxygenation and high perfusion
- Q4 (White) – Low oxygenation and low perfusion

To generate the VFI maps, individual StO_2_ and power Doppler images were first independently thresholded using empirically determined cutoffs: 60% for oxygen saturation and 70% of the maximum PDI signal intensity for perfusion. These thresholds were selected based on physiological relevance and visual inspection across multiple datasets to distinguish functionally meaningful regions. To ensure consistency and reproducibility, representative pre-treatment images from all experimental groups were randomly sampled, and an unsupervised two-group clustering analysis was performed to determine optimal threshold values for both modalities. The tumor region of interest was then segmented, and each pixel was assigned to one of the four VFI categories based on its local StO_2_ and Doppler intensity values. The resulting color-coded VFI maps, as shown in Figure 5, provide a spatially intuitive representation of heterogeneous tumor physiology, allowing visualization of how PDT perturbs both oxygenation and vascular integrity over time. For quantitative analysis, the number of pixels within each quadrant was computed and normalized to the total number of tumor pixels, resulting in fractional VFI metrics (VFI_Q1, VFI_Q2, VFI_Q3, VFI_Q4). These fractions were used to monitor temporal changes in the vascular functional landscape of the tumor post-treatment and to assess therapeutic efficacy.

### Histology

Tumors were harvested at designated endpoints and embedded in optimal cutting temperature (OCT) compound for cryosectioning. Serial sections, 10□µm thick, were obtained using a cryotome, maintaining the same imaging plane orientation as that used in the corresponding PAI datasets. For histological analysis, every alternate section was stained with hematoxylin and eosin (H&E) following standard protocols previously described [41]. These slides were used to assess overall tumor architecture and morphology. Adjacent sections were processed for immunofluorescence (IF) staining to evaluate vascular integrity and hypoxia.

For immunofluorescence staining, tumor sections were incubated overnight at 4□°C with primary antibodies targeting vasculature and hypoxia: anti-mouse CD31 (Goat polyclonal IgG, R&D Systems, Cat: AF3628; 1:5 dilution) and anti-pimonidazole (conjugated Rat monoclonal IgG1, clone 11.23.22.r, Hypoxyprobe, Cat: Red 549 Mab; 1:25 dilution). The following day, slides were washed with PBS and incubated with a secondary antibody, NorthernLights™ 637 Anti-goat IgG (R&D Systems, Cat: NL002; 1:50 dilution), for 2 hours at room temperature. Nuclear counterstaining was performed using DAPI-containing antifade mountant (SlowFade™ Gold with DAPI, Thermo Scientific, Cat: S36939), as described in Langley et al., before cover slipping and imaging the slide [57].

Fluorescence images were acquired using the EVOS M7000 imaging system equipped with LED-based filter cubes suitable for DAPI, red, and far-red fluorescence channels. H&E and IF images were further processed in FIJI (ImageJ) to enhance clarity for visualization. For IF images, minor adjustments to brightness and contrast were applied to emphasize signal distribution. Non-relevant areas, such as skin and background, were cropped for clarity.

## Statistics

All statistical analyses were performed using GraphPad Prism (La Jolla, CA). A two-way ANOVA with Tukey’s post hoc test was used to evaluate the effects of treatment (NT, 50 J/cm^2^, and 100 J/cm^2^) and time (pre-treatment, 1 hour, 6 hours, and 24 hours) on % StO_2_, % PDI, and all the Vascular fractional Index (VFI) parameters. Treatment was considered a between-subjects factor, while time was treated as a within-subjects factor. All results are reported as mean ± standard error of the mean (SEM), and a p-value < 0.05 was considered statistically significant.

## Acknowledgments

The authors would also like to acknowledge the members of the integrated Biofunctional Imaging and Therapeutics laboratory (Allison Sweeney, Macy Halim, Rebecca Moriarty, Brooke Bednarke, Avijit Paul and Andrew Langley) for their help and support

## Author contributions

Conceptualization: SM, PK, DMS

Methodology: DMS, LW, RTS

Investigation: DMS, LW

Visualization: DMS

Supervision: SM

Writing—original draft: DMS

Writing—review & editing: LW, RTS, PK, SM

## Data and materials availability

All data are available in the main text or the supplementary materials.

